# Limited adaptive evolution of *Staphylococcus aureus* during transition from colonization to invasive infection

**DOI:** 10.1101/2021.10.25.465068

**Authors:** Anna K. Räz, Federica Andreoni, Mathilde Boumasmoud, Judith Bergada-Pijuan, Tiziano A. Schweizer, Srikanth Mairpady Shambat, Barbara Hasse, Annelies S. Zinkernagel, Silvio D. Brugger

**Affiliations:** Department of Infectious Diseases and Hospital Epidemiology, University Hospital Zurich, University of Zurich, Zurich, Switzerland

**Keywords:** *Staphylococcus aureus*, colonization, invasive infection, transition, barrier breach

## Abstract

*Staphylococcus aureus* carriage is a risk factor for invasive infections. Unique genetic elements favoring the transition from colonizing to invasive phenotype have not yet been identified and phenotypic traits are understudied. We therefore assessed pheno- and genotypic profiles of eleven *S. aureus* isolate pairs sampled from colonized patients simultaneously suffering from invasive *S. aureus* infections. Ten out of 11 isolate pairs presented the same *spa* and MLST suggesting colonization as origin for the invasive infection. Systematic analysis of colonizing and invasive isolate pairs showed similar adherence, hemolysis and reproductive fitness properties, minimal genetic differences, identical antibiotic tolerance and bacterial virulence in *Galleria mellonella*. Our results provide insights into the similar phenotype associated with limited adaptive genetic evolution between the colonizing and invasive isolates. Disruption of the physical barriers mucosa or skin were identified in the majority of patients further emphasizing colonization as a major risk factor for invasive disease.

**Importance:** *S. aureus* is a major pathogen of humans causing a wide range of disease. The failure to develop a vaccine and antibiotic warrant the exploration of novel treatment strategies. Asymptomatic colonization of the human nasal passages is a major risk factor for invasive disease and decolonization procedures have shown efficacy in preventing invasive infections. However, the transition of *S. aureus* from a benign colonizer of the nasal passages to a major pathogen is not well understood and host as well as bacterial properties have been discussed as being relevant for this behavioral change. We conducted a thorough investigation of patient-derived strain pairs reflecting colonizing and invasive isolates in a given patient. Although we identified limited genetic adaptation in certain strains as well as slight differences in adherence capacity of colonizing and invasive isolates, our work suggests that barrier breaches are a key event in the disease continuum of *S. aureus*.

## Introduction

*Staphylococcus aureus* is a frequent asymptomatic colonizer of the human skin and mucosa and is identified as part of the normal human microbial flora (1). *S. aureus* carriage can be persistent (20%) or intermittent (60%) (2). Nevertheless, colonization puts the individual at increased risk for developing invasive infections, ranging from simple skin to life-threatening infections (1, 3). *S. aureus* is therefore classified as a pathobiont reflecting this dual behavior of a colonizer and a pathogen and the corresponding damage-response continuum, capable of infecting healthy as well as immunocompromised individuals (4, 5).

In industrialized countries the annual incidence rate for *S. aureus* blood stream infection is 26.1 per 100’000 inhabitants with a case-fatality rate of around 25% (6). Treatment of *S. aureus* infections faces various obstacles, such as antibiotics resistance and bacterial persistence (7). Given the burden of *S. aureus* disease, different prevention strategies are under investigation. Decolonization has proven efficient in preventing nosocomial *S. aureus* infections among carriers (3, 8)

The exact circumstances under which a colonizing *S. aureus* strain becomes invasive and causing host damage are incompletely understood. Invasive *S. aureus* infections most probably originate from the bacteria colonizing the nose and throat. A breach of the protective physical barriers of the skin and mucosa allows the bacteria to enter the bloodstream and spread to other body sites (9). Information supporting this hypothesis comes from studies comparing invasive and colonizing *S. aureus* strains isolated from the same patient where over 80% of isolate pairs were identical, as determined by pulsed-field gel electrophoresis (10, 11). Research carried out in this field compared genotypes, virulence gene expression of colonizing and invasive *S. aureus* strains, as well as factors influencing the transition (12–17). However, to our knowledge a pheno- and genotypic assessment of strain adaptation during transition from a colonizing to an invasive behavior by matching colonizing and invasive isolates derived from the same patient is still missing. The aim of this study was therefore to analyze the genetic traits, virulence, reproductive fitness and antibiotic tolerance capacity of corresponding colonizing and invasive *S. aureus* strains, isolated from the same patient and referred to as isolate pairs.

## Materials and Methods

### Patient recruitment

The study was conducted as part of the BacVivo “Bacterial pathogen properties in patient samples” study, a single-center observational study conducted at the University Hospital Zurich. This study was approved by local authorities (cantonal ethics committee, canton of Zurich, ethical application number: BASEC 2017-02225). Patients diagnosed with an invasive *S. aureus* infection were screened for the presence of colonizing *S. aureus* in the nose and groin during their hospital stay. Patients with colonizing *S. aureus* were included in this analysis (Tab.1). All patients included in this study signed an informed consent. Healthy volunteers’ blood was collected under the protocol BASEC 2019-01735, approved by the cantonal ethics committee, Canton of Zurich.

### *S. aureus* clinical isolates and processing of clinical samples

*S. aureus* strains were isolated from either blood, tissue, nose or groin (Tab.2). Blood was cultured in BacT/Alert FA bottles (Biomérieux) at 37°C to positivity or up to 6 days. Tissue samples were homogenized using a mortar. Samples from homogenized tissues and swabs were cultured in thioglycolate broth to positivity. Antibiotic susceptibility testing (Tab.S1) was carried out on Müller Hinton agar by disk-diffusion test, EUCAST breakpoints were used for interpretation of the results (18).

### *Spa* typing

*spa* typing was carried out as previously described (19). See supplemental material for a detailed description.

### Whole Genome Sequencing and Comparative Genomics

DNA libraries were prepared with the QIASeq FX DNA library kit (Qiagen) and sequenced with the MiSeq Reagent Kit V2 (Illumina) on a MiSeq instrument. The 150 bp paired end read files were cleared from Illumina adaptors and low-quality reads with Trimmomatic v.0.39 (20). De novo-assemblies were built with SPAdes v.3.10 with the --careful command and default k-mer size (21). The contigs with less than 2-fold average k-mer coverage and shorter than 200bp were discarded. For multi-locus sequence type (ST) determination, the assemblies were scanned against traditional PubMLST typing schemes (22) with MLST v 2.7.6 (23). Annotation of our assemblies and the assemblies of selected reference genomes (Tab.S2) was performed with Prokka v.1.13 (24). The GFF files generated by Prokka were used as input to Roary v.3.12 for pangenome construction (25). The command -e --mafft was used to generate a multi-fasta alignment of the core genes. Fasttree v.2.1.10 (26) was used to build a maximum likelihood tree, with the default Juke Cantor model of nucleotide evolution. Variant calling between pairs of isolates was performed with Snippy v.4.4.3 (27), by aligning the matching pairs’ trimmed reads to the *de novo* assembly of one of the isolates as well as to the .gbk file of the closest reference to assess the effects of variants. Illumina paired-end reads of the 22 isolates are available through the European Nucleotide Archive project PRJEB47806.

### Hemolysis assay

Hemolysis capacity of the various *S. aureus* strains was determined as previously described (28). See supplemental material for a detailed description.

### Adherence assay

Adherence of the *S. aureus* clinical isolates to human lung alveolar basal epithelial cells (A549) was tested as previously described (29).

### Quantitative Fitness Analysis (QFA)

To compare the reproductive fitness of the clinical isolates, Quantitative Fitness Analysis (QFA) was carried out as previously described (30)

### Minimal inhibitory concentration

The minimal inhibitory concentration (MIC) for the antibiotics ciprofloxacin, clindamycin and flucloxacillin was tested according to the broth microdilution method, following the recommendations of the “European Committee on Antimicrobial Susceptibility Testing of the European Society of Clinical Microbiology and Infectious Disease” (31). MICs were tested in pH 7.4 medium (35 ml DMEM + 10% FBS +1% L-glutamine mixed with 12.62 ml H2O and 2.38 ml of 1M HEPES buffer).

### Persister assay

To assess the fraction of persisters in the *S. aureus* clinical isolates population, an assay based on colony size distribution and bacterial survival to antibiotics upon low pH stress was conducted as previously described (32, 33). Images were analyzed with ColTapp (34).

### *Galleria mellonella* virulence assay

To study *S. aureus* virulence in vivo, the non-mammalian model system *Galleria mellonella* was used (35). 6 *Galleria mellonella* per group were injected with 10^4^ log phase S. aureus CFUs A control group was injected with 10μl of PBS and a second control group of non-injected *Galleria mellonella* was also used. The larvae were subsequently incubated at 37°C and assessed at 24, 48 and 72 hours post infection. See supplemental material for a detailed description.

### Statistics

Statistical analyses were carried out using GraphPad Prism (Version 8, GraphPad Software, Inc.). To test for differences in cell adherence, hemolysis and percentage of the subpopulation persisters between isolate pairs and between invasive and colonizing strains, the Wilcoxon matched-pairs signed rank test or paired t-test were used. To compare *G. mellonella* survival curves a log-rank test was conducted. P values of < 0.05, two-tailed, were considered statistically significant.

## Results

### Patients cohort and clinical isolates

Thirty-eight patients suffering from distinct invasive *S. aureus* infections, were enrolled in the BacVivo study between January and November 2019. Among these, 11 tested positive for *S. aureus* colonization in the nose or groin and were selected for this project (Tab.1). Patients showed a variety of different *S. aureus* infections ranging from surgical wound infection to extensive skin infections and bacteremia. The patient cohort consisted of seven male and four female patients, age ranging between 29 to 86 years. In seven patients a barrier breach was associated with infection.

### Antibiotic susceptibility testing

Two out of the 22 strains, both collected from patient 9, were methicillin-resistant *Staphylococcus aureus* (MRSA). Antibiotic susceptibility testing results are listed in Tab.2 and Tab.S1.

### Strains *spa* and MLST characterization

In order to assess potential clonality of the colonizing and invasive *S. aureus* isolates, all clinical isolates included in this study were initially *spa*-typed. Twelve different *spa* types were found (Tab.2). In 10 out of 11 cases, the invasive and the colonizing *S. aureus* strains showed the same *spa* and multi locus sequence typing (MLST) type and among the matching pairs one new *spa* type (t19435) and two new MLST profiles (ST-6099 and ST-6100) were found and submitted to PubMLST (22).

### Whole genome sequencing reveals minimal genomic evolution during transition from colonization to invasion

To investigate genetic differences as a measure of genetic evolution during transition from colonization to invasive disease, whole genome sequencing of all isolates was performed. The diversity of the isolates from patient 8 was confirmed, as illustrated by their distance on the core genome-based phylogenetic tree (Fig.S1). All other pairs of isolates clustered together and a reference strain closely related to each pair was chosen for comparative genomics (Fig.S1, Tab.S2).

None of the ten pairs of isolates with matching *spa*-type and MLST displayed gene content differences. At the core genome level, 40% of them (4/10), namely CI3990-CI3984 (patient 7), CI4130-CI4099 (patient 9), CI4097-CI4090 (patient 10) and CI4152-CI4116 (patient 11), were found to be identical. The remaining six pairs displayed between 1 and 26 core genome variants (insertion, deletions and single nucleotide polymorphisms) (Tab.S3). Of these six pairs, two, CI2670-CI2647 (patient 5) and CI3203-CI3203 (patient 6) displayed only synonymous or outside-from-coding-sequences mutations. This was the case for 64% (51/80) of the total variants identified. The 29 non-synonymous variants, affecting CI1418-CI1818 (patient 1), CI2016-CI2025 (patient 2), CI2056-CI2050 (patient 3), CI2182-CI2327 (patient 4) are shown in Tab.3. Most were missense variants (n=23) and the remaining were one in-frame deletion, four frameshift variants and one premature stop.

Among other mutations, a missense variant (Tyr95His) of *agrA* distinguished CI1818 from CI1814. Within-host mutations of this master regulator gene of *S. aureus* virulence have been previously described and are expected to be phenotypically relevant. In addition, a frameshift variant (Gly4629fs) of Ebh, the extracellular matrix-binding protein might result in altered adherence capacity, complement resistance and pathogenesis during infection (36–39). Among the variants distinguishing CI2016 from CI2025, a missense variant of gdpP could be phenotypically relevant. Finally, a missense variant (Arg244His) of MutM, a protein involved in DNA mismatch repair, distinguished CI2182 from CI2327. An increased mutation rate would be a plausible explanation for the substantial diversification of this strain as compared to others: the 26 variants between its two isolates were the highest genetic distance across pairs. In conclusion, the clonal origin of the isolate pairs was confirmed and different extent of diversification could be observed in different patients.

### Comparable hemolysis, adherence properties and reproductive fitness of colonizing and invasive *S. aureus* isolate pairs

We next characterized phenotypically the pairs of isogenic isolates. The hemolytic capability of the *S. aureus* clinical isolates was tested. However, no significant difference was found across strains or between colonizing and invasive isolates (2.697 AU vs 2.818 AU, p=0.5527) (Fig.1A). Since adherence to host cells is the first step leading to invasion and spread of a bacterial infection, we compared the adhesion capacity of invasive versus colonizing isolates. Although adherence to the lung epithelial cell line A549 showed significant difference between colonizing and invasive isolates (4.199% vs. 3.462%, p=0.0244), the biological relevance of this difference is debatable (Fig.1B). To compare the growth characteristics and reproductive fitness, quantitative fitness analysis (QFA) was performed. Colonizing versus invasive isolates showed no difference in reproductive fitness (0.001 vs. 0.001, mean difference 1.28e-6, p=0.9766) (Fig.1C).

**Figure 1.**
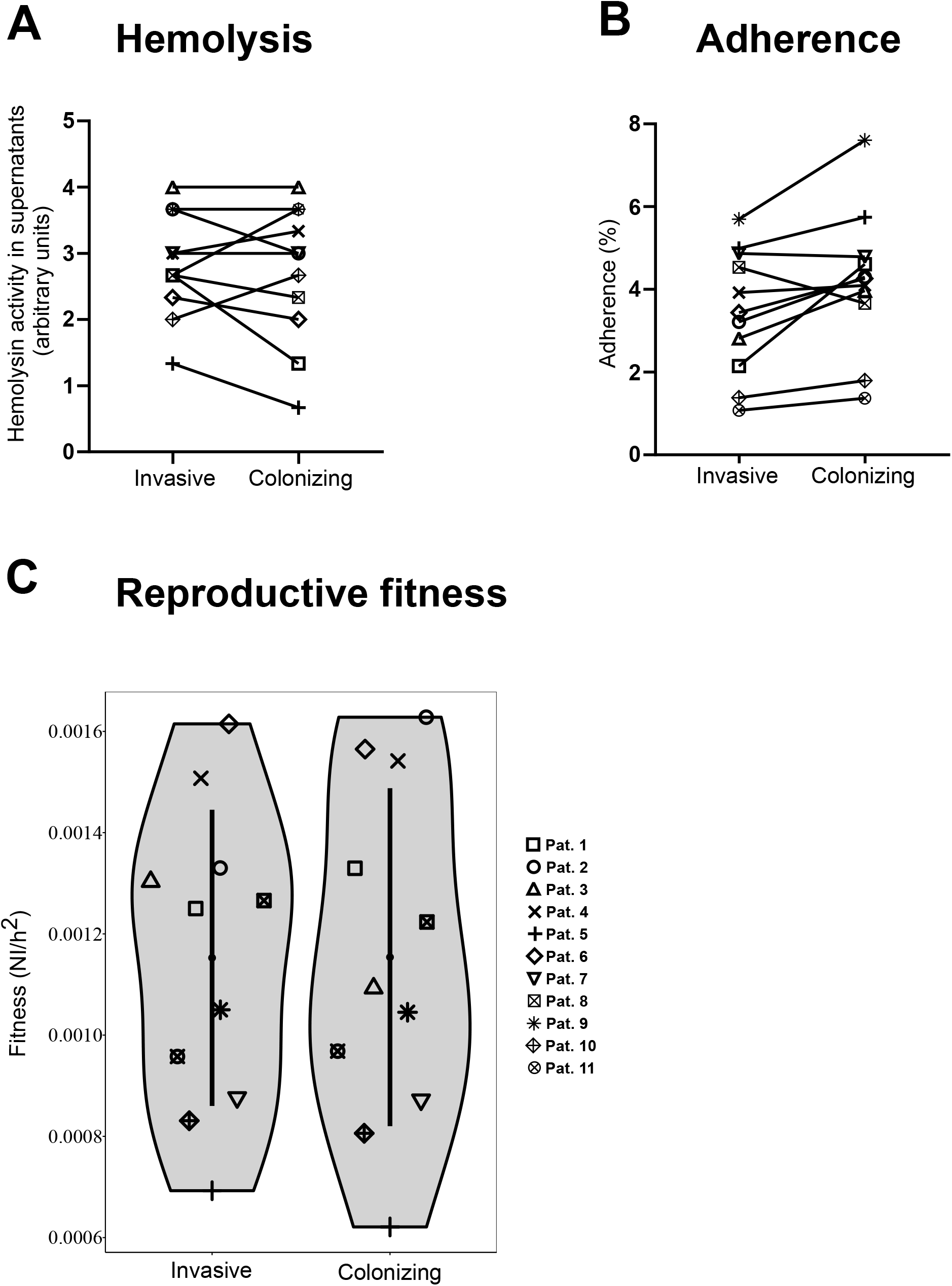
Comparable hemolysis, adherence properties and reproductive fitness of colonizing and invasive *S. aureus* strains and strains pairs. **A)** Hemolysis activity detected in *S. aureus* clinical isolates supernatants. 1:2 serial dilutions of supernatants derived from OD_600_ 6 overnight cultures of the strains of interest were incubated with human erythrocytes for 1h at 37°C followed by 30min at 4°C. Absorbance at 415nm was measured to detect hemolysis and the last dilution displaying activity was plotted on the graph. The graph displays pooled data from three independent experiments. **B)** Adherence of *S. aureus clinical* isolates to human adenocarcinoma human lung alveolar basal epithelial cells (A549). Mid-logarithmic growth-phase bacteria were added to A549 cells a multiplicity of infection (MOI) of 10. After incubation at 37°C in 5% CO_2_ for 30 min the cells were washed with PBS, lysed with PBS + 0.02% triton-X100 and adhering bacteria were enumerated by plating serial dilutions of this suspension. The graph displays pooled results from three independent experiments. **C)** Reproductive fitness of invasive and colonizing S. aureus clinical isolates. Reproductive fitness was calculated, according to the Gompertz mathematical model, as the ratio between maximum growth-rate (MGR) and time to reach MGR (T_max_) displayed by the various isolates. NI/h^2^ = normalized intensity per square hour; Pat = patient.

### Persisters formation and colony size distribution do not vary between colonizing and invasive *S. aureus* isolates

In order to evaluate the phenotypic plasticity of the isolates upon environmental stress, we assessed their ability to form persisters upon acidic stress in vitro. After pH stress exposure the clinical isolates were challenged with antibiotics. Generally the different strains responded in a very similar manner to pH stress followed by antibiotic challenge (Fig.2A).

**Figure 2.**
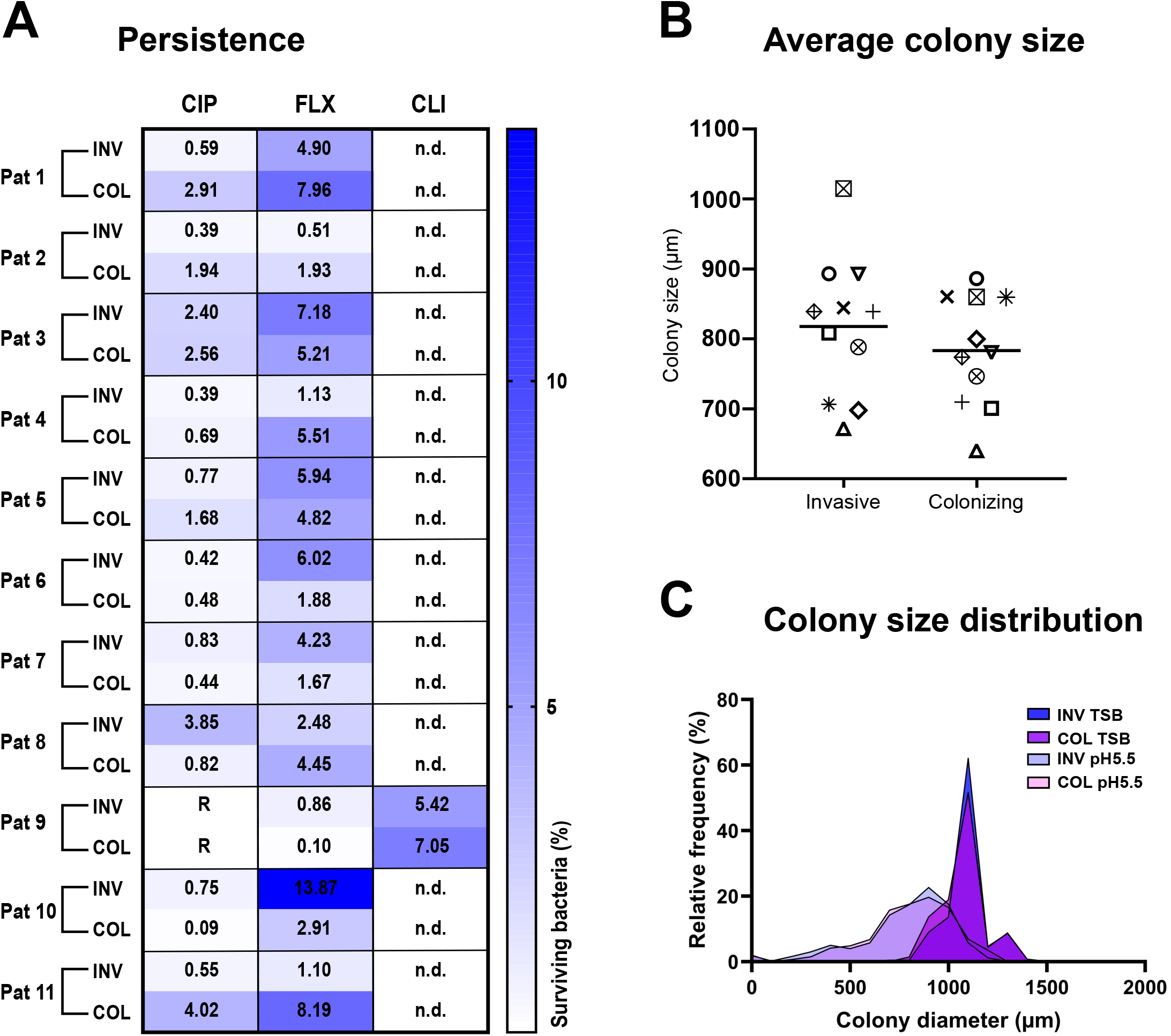
Persisters formation and colony size distribution do not vary between colonizing and invasive *S. aureus* strains and strains pairs. **A)** Persisters formation was assessed on *S. aureus* strains after incubation in pH 5.5 medium for 72h at 37°C in 5% CO_2_ followed by incubation for 24h in pH 7.4 medium in the presence of absence of 40 times the minimal inhibitory concentration (MIC) of either ciprofloxacin (CIP), flucloxacillin (FLX) or, if ciprofloxacin resistant, clindamycin (CLI). After 24h the antibiotics were washed out and the surviving bacteria were plated on TSB plates and counted. The figure shows the surviving bacteria as a percentage of the inoculum. At least 4 biological replicates were carried out per strain. **B)** Average colony size was assessed for invasive and colonizing strains by measuring the colony size of the *S. aureus* clinical isolates after 72h of pH 5.5 stress and subsequent incubation at 37°C for 24 hours on blood agar plates. **C)** colony size distribution was assessed for invasive and colonizing strains by measuring the colony size of the *S. aureus* clinical isolates after 72h of pH 5.5 stress or 72h of growth in TSB and subsequent incubation at 37°C for 24 hours on blood agar plates. A minimum of 3 biological replicates were carried out per strain. Pat=patient, INV=invasive, COL=colonizing, R=resistant, n.d.=not determined.

The persisters triggered by pH stress need some time to exit their dormancy state, which results in non-stable small colonies (nsSCs) upon plating on solid nutrient medium. nsSCs can be detected from a bacterial population and are an index of the presence of persisters (28, 32). Hence, we also characterized the colony size heterogeneity of bacteria upon pH challenge as compared to bacteria grown in rich medium (TSB). No significant differences in average colony size were found across colonizing and invasive isolates (817.7 um vs 783.3 um, p=0.5190) or between isolate pairs after pH stress (Fig.2B). A general shift in colony size distributions, with long tails of small colonies, was observed upon pH stress as compared to growth in rich medium (Fig.2C). This was independent of the strains’ origin of being either colonizing or invasive.

### Similar virulence of colonizing and invasive *S. aureus* isolates in vivo

*G. mellonella* larvae were used to investigate the virulence of the *S. aureus* clinical isolates in vivo. Larvae injected with colonizing or invasive *S. aureus* isolates showed a significantly decreased survival as compared to PBS-injected ones (invasive isolates: p= 0.0023; colonizing isolates p= 0.0002) and no difference between colonizing and invasive isolates was found (p=0.3541) (Fig.3). There was no significant difference between the survival of PBS-injected and non-injected larvae.

**Figure 3.**
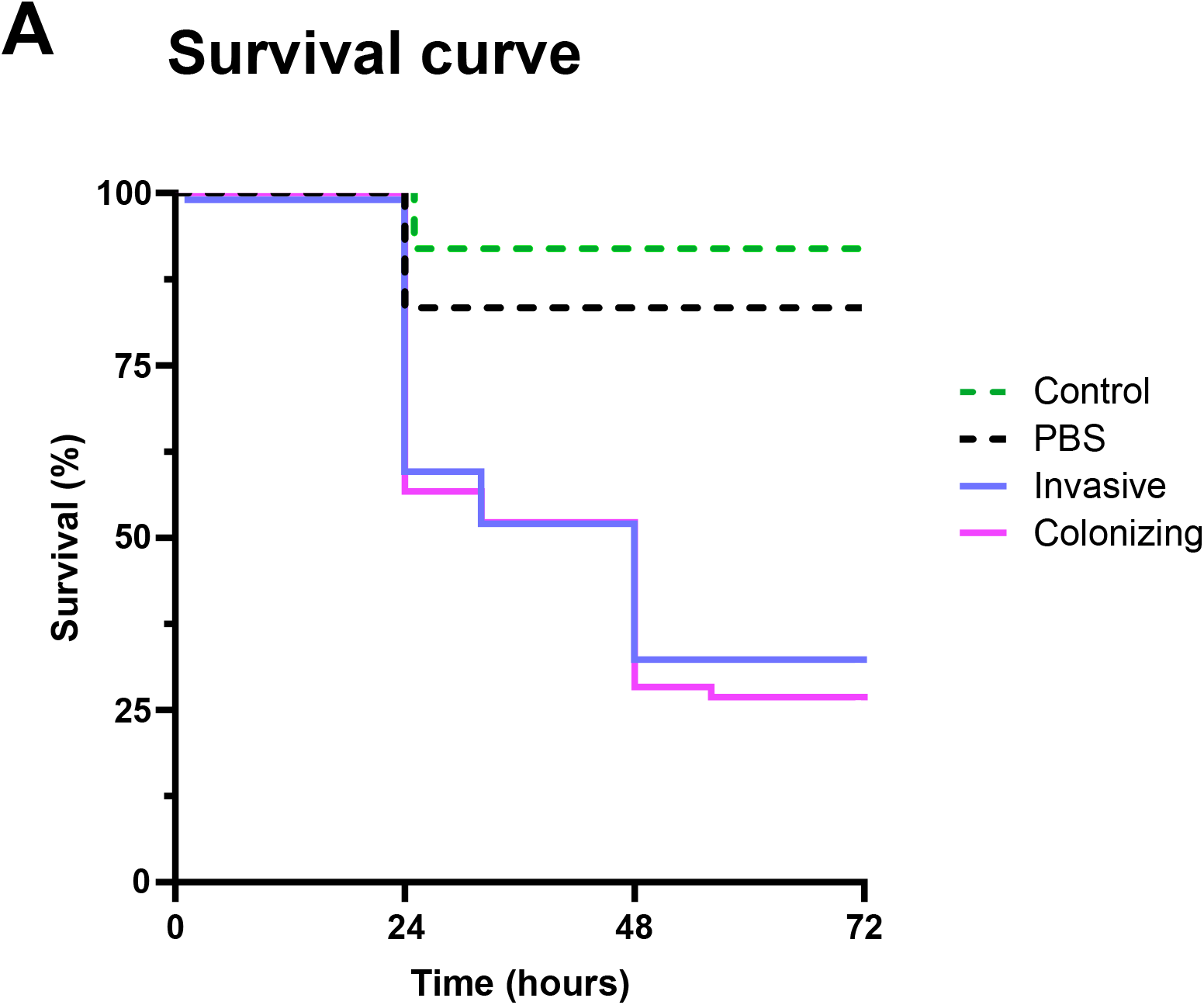
Effect of *S. aureus* clinical isolate strains on the viability of *Galleria mellonella* larvae over 72 h. *Galleria mellonella* were infected with 10^4^ colony forming units in 10μl PBS. Afterwards, *Galleria mellonella* were incubated at 37°C and the viability was assessed over a 72 hours period. Each test group contained 6 *Galleria mellonella* and the graph shows the results of one experiment. Statistical differences between the survival curves were assessed using the Log-rank test.

## Discussion

We characterized 11 colonizing and invasive *S.aureus* pairs from patients with invasive *S.aureus* infections and found a similar phenotype associated with limited adaptive genetic evolution between the colonizing and the invasive isolates. Breaching of the skin or mucosa was found in most of the patients underlining the observation that colonization is a major risk factor for *S. aureus* invasive infections. So far various studies have assessed genotypic characterization but failed to pinpoint traits responsible for the transition from colonizing to invasive behavior. Aiming to assess whether phenotypic properties would influence the invasion characteristics of the isolated strains, extensive analysis of virulence traits of *S. aureus* isolate pairs was carried out using well established *in vitro* and *in vivo* models coupled with techniques for the detection of *S. aureus* fitness and persisters. Despite the broadness of approaches, no significant differences in virulence or environmental stress adaptation could be detected, mirroring results obtained previously (16), calling for a more in-depth analysis of the role of host factors in the transition between colonizing and invasive phenotypes.

In a first step, we screened the invasive and colonizing clinical isolates for their genetic background and antibiotic susceptibility spectrum. We found that the isolate pairs shared, in 10 out of 11 cases, the same *spa-type* and MLST, thus strengthening the hypothesis that colonizing *S. aureus* strains are the reservoir from which invasive infections originate. The genetic concordance between colonizing and invasive strains isolated from the same individual was shown before (10, 11, 40), confirming a common pattern.

To compare the virulence of our panel of clinical isolates we investigated the role of two key virulence factors in the pathology of *S. aureus* infections, hemolysis and adherence to eukaryotic cells. The pore-forming toxin α-hemolysin, also known as α-toxin, is a main factor of *S. aureus* influencing hemolysis (41, 42) and its activity was measured for colonizing and invasive clinical isolates. Our results show that colonizing strains do not differ from invasive strains in their capacity to lyse red blood cells, in the conditions tested using our experimental settings. The ability to adhere to and invade epithelial cells plays a major role in *S. aureus* colonization, biofilm formation as well as early manifestation of *S. aureus* infection and was previously shown to vary among different *S. aureus* strains (43). Furthermore, irrespective of the location, adherence increases the chance of the bacterium to breach defensive barriers (44). We tested adherence of the *S. aureus* clinical isolates to lung epithelial cells. A significantly higher adherence was observed for colonizing strains although the biological relevance of this difference remains uncertain. Comparison of growth dynamics of the various clinical isolates allowed screening for changes in reproductive fitness possibly linked to the transition towards an invasive phenotype. Overall, we found a large variation in reproductive fitness among the *S. aureus* isolates. However, the fact that we didn’t observe differences between isolate pairs or between colonizing vs invasive isolates indicates that reproductive fitness does not play a role in the transition from colonizing to invasive phenotype.

Survival in the host is a key factor that allows bacterial spread during an invasive infection. Exposure of bacteria to environmental stress, such as low pH or residing in the host, leads to formation of a sub-population of persister cells (28, 32, 45), characterized by a growth-arrest phenotype that allows them to survive both host defense mechanisms and antibiotic treatment targeting active cellular processes. We hypothesized that invasive isolates may display a higher degree of persister cell formation due to the exposure to host environmental stressors such as pH, antibiotics and immune cells. We therefore examined the capability of colonizing and invasive *S. aureus* isolates to form a sub-population of persister cells upon pH stress and subsequent antibiotic exposure. Survival of invasive and colonizing clinical isolates did not differ, with all strains displaying a very similar survival profile. This suggests that the ability to form persisters upon pH stress is similar for both colonizing and invasive isolates. Persisters, as a result of their slow-growth phenotype, generate phenotypically heterogeneous nsSC (32). As an estimate of the presence of a persister population, we also compared the average colony size and colony size distribution of colonizing and invasive clinical isolates after pH stress. No difference in average colony size was found, indicating a similar capacity to form a persister population in both sets of isolates. Colony size distribution plots, in which we compared the colony size of strains grown in nutrient-rich medium and neutral pH with the colony size of strains grown in nutrient-poor medium under pH stress conditions, indicate the presence of a consistent subpopulation of persisters in all clinical isolates.

Next, we tested the virulence of the clinical isolates *in vivo*, using the larvae of *G. mellonella* as a model organism (46). Colonizing and invasive clinical isolates were compared for their ability to infect and kill the larvae. No survival differences were found, neither between the two groups of isolates, nor between strains pairs.

Whole genome sequence analysis confirmed the absence of specific determinants causing the transition to a more virulent phenotype, in line with previous work showing similar gene content between invasive and colonizing isolates (47). Four out of the 10 isolate pairs were genetically identical, while few mutations distinguished the remaining 6 couples. Of these, none was recurring in several pairs. Previous works carried out on *S. aureus* isolates also showed equal occurrence of virulence genes among *S. aureus* colonizing and invasive isolates mirroring our results on larger samples’ cohorts (14, 16, 47, 48). However, some isolates contained mutations that might reflect adaption and other groups have reported mutations associated with the transition from colonization to invasive disease or during invasive disease (12, 49). Abu Othman et al also tested virulence genes expression of *S. aureus* clinical isolates during growth in the presence of human serum, to better mimic the host conditions (17). A high variation in gene expression among different clinical isolates was observed and the expression of the collagen adhesion-encoding gene *cna* was upregulated in colonizing isolates while γ-hemolysin Hlg was upregulated in invasive isolates confirming that the environmental conditions can influence strains behavior (17). Another interesting study showed the importance of environmental changes for virulence determinants expression by comparing *S. aureus* virulence factors expression in animal models of colonization, bacteremia and endocarditis, pinpointing increased expression of a set of genes as important for the transition from colonizing to invasive behavior (50).

In conclusion, we examined several genetic and phenotypic characteristics that confer virulence to *S. aureus*, aiming at elucidating main players involved in the transition from colonizing to invasive behavior. The absence of significant differences in virulence between invasive and colonizing strains we observed in this study does not reflect the disease progression observed in patients, where seemingly innocuous colonizing strains invaded and spread in the host, causing severe manifestations. The lack of differences between colonizing and invasive isolates are attributable to environmental and host factors such as breaching of the physical barrier. This is underlined by previous work which also suggests the likelihood of a host factors-mediated transition (47, 51). As colonization is the preponderant risk factor for invasive staphylococcal disease and given the limited success of decolonization procedures, novel approaches such as decolonization of *S. aureus* with transplantation of beneficial bacterial commensals might be an attractive and efficient way to prevent invasive diseases.

## Acknowledgments

We would like to thank Sebastian Herren and Kim Alan Röthlin for their support with strain acquisition.

## Author Contributions

AKR: Experimental design, acquisition, analysis and interpretation of data, writing of the manuscript

FA: Experimental design, acquisition, analysis and interpretation of data, writing of the manuscript

MB: Analysis and interpretation of data, writing of the manuscript

JBP: Analysis and interpretation of data

TS: Experimental design, data analysis and critical reading of the manuscript

SMS: Experimental design, data analysis and critical reading of the manuscript

BH: Conceptualization, funding and critical reading of the manuscript

ASZ: Conceptualization, funding and critical reading of the manuscript

SDB: Conceptualization, experimental design, acquisition and interpretation of data, funding, writing of the manuscript

---

**Table 1.**
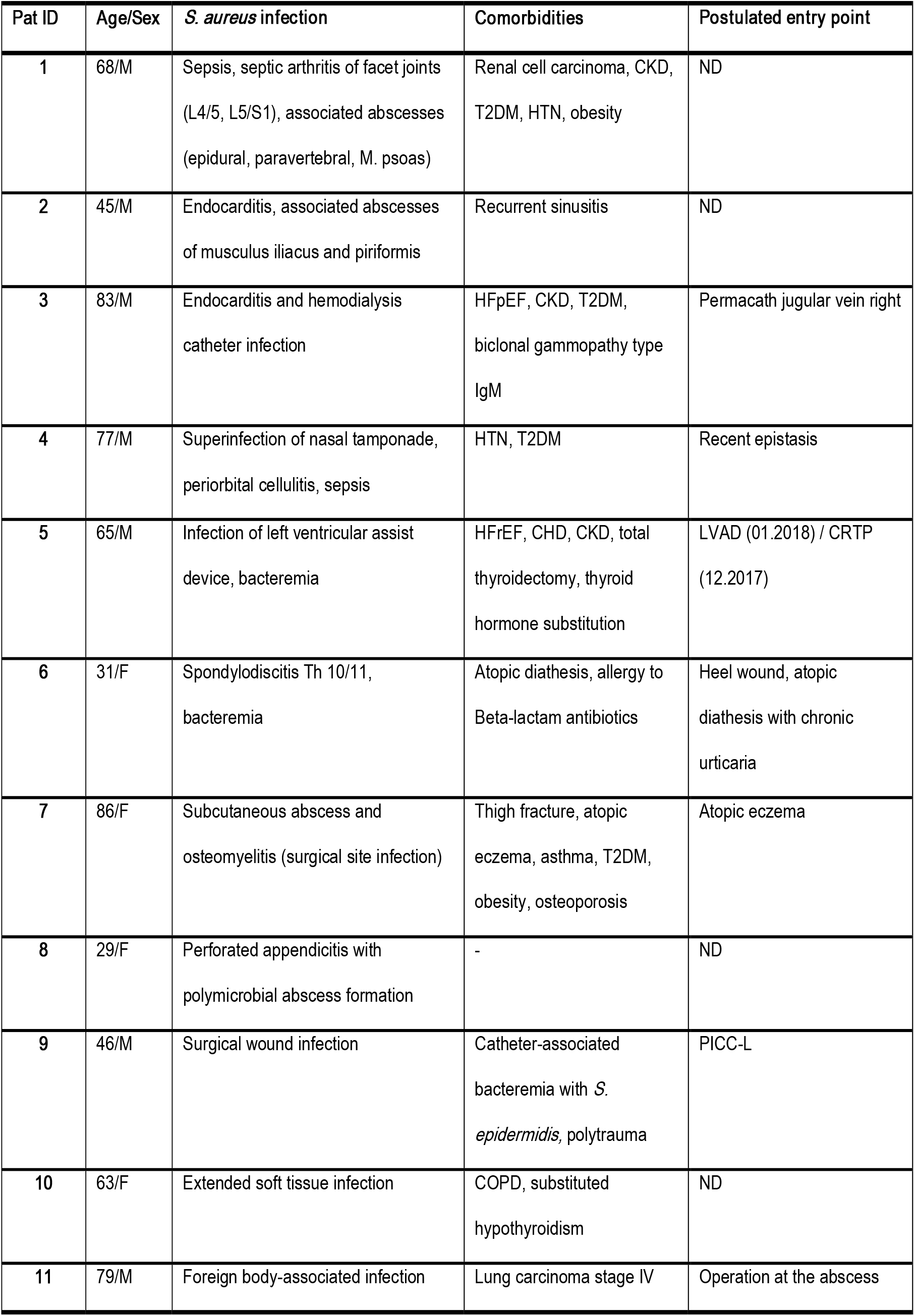
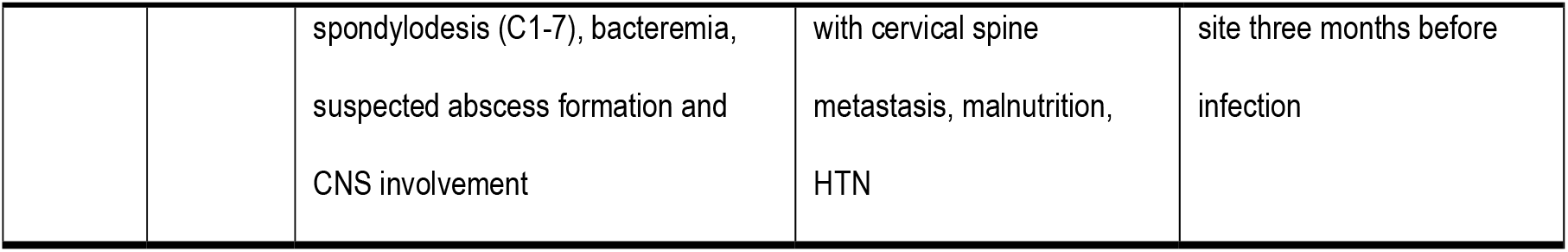
Patient characteristics. Patients’ demographic data, infection type and relevant comorbidities. M=male, F=female, CNS=central nervous system, CKD=chronic kidney disease, T2DM=type-2 diabetes mellitus, HTN=hypertension, HFpEF=heart failure with preserved ejection fraction, HFrEF= heart failure with reduced ejection fraction, CHD=coronary heart disease, COPD=chronic obstructive pulmonary disease, LVAD= Left Ventricular Assist Device, CRTP= Cardio Resynchronisation Therapy Pacemaker, PICC-L= Periferally Inserted Central Catheter-Line, ND=not determined.

**Table 2.**
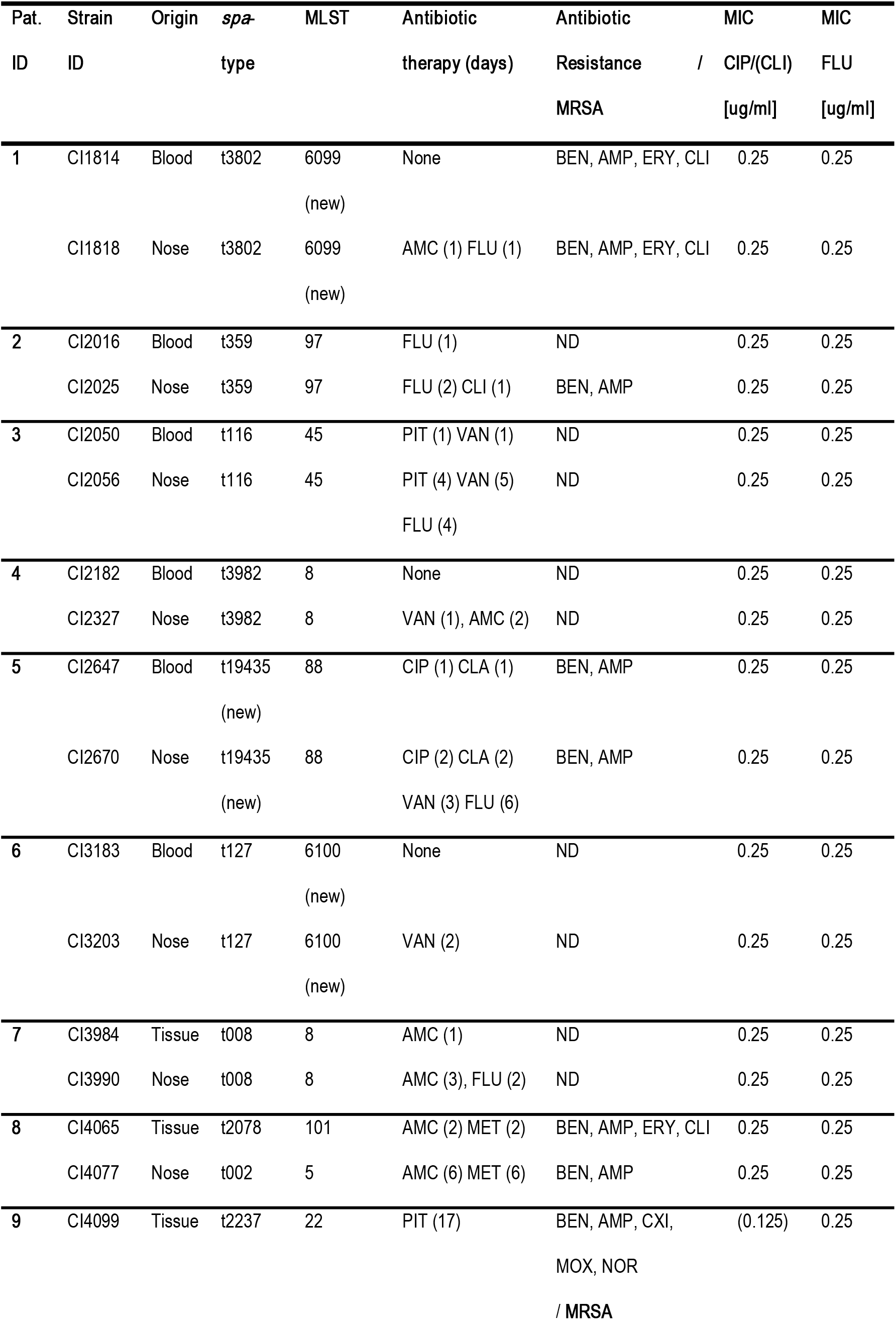
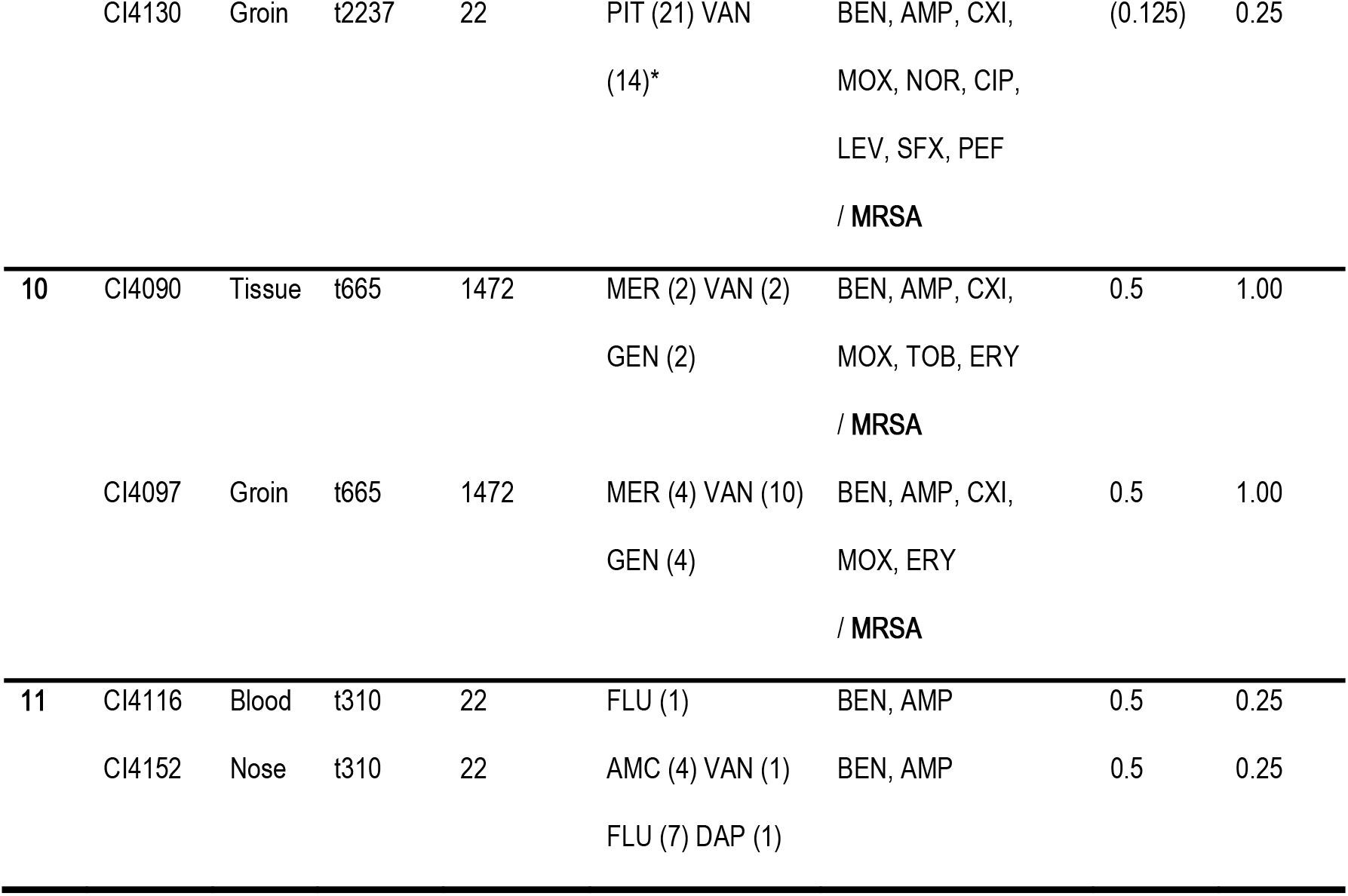
Clinical isolates information. All strains information, including strain ID, origin, *spa* (staphylococcal protein A)-type and multilocus sequence (MLST)-type, antimicrobial therapy before isolation (in days), resistance to tested antibiotics, classification as methicillin-resistant *Staphylococcus aureus* (MRSA) and minimal inhibitory concentration (MIC) for Ciprofloxacin (CIP), Clindamycin (CLI) and Flucloxacillin (FLU) can be found in table 1 (18). ND=none-detected (no antibiotic resistance detected for the tested antibiotics), AMC=Amoxicillin-clavulanate, AMP=Ampicillin, BEN=Benzylpenicillin, CLA=Clarithromycin, CXI=Cefoxitin, DAP=Daptomycin.

**Table 3.**
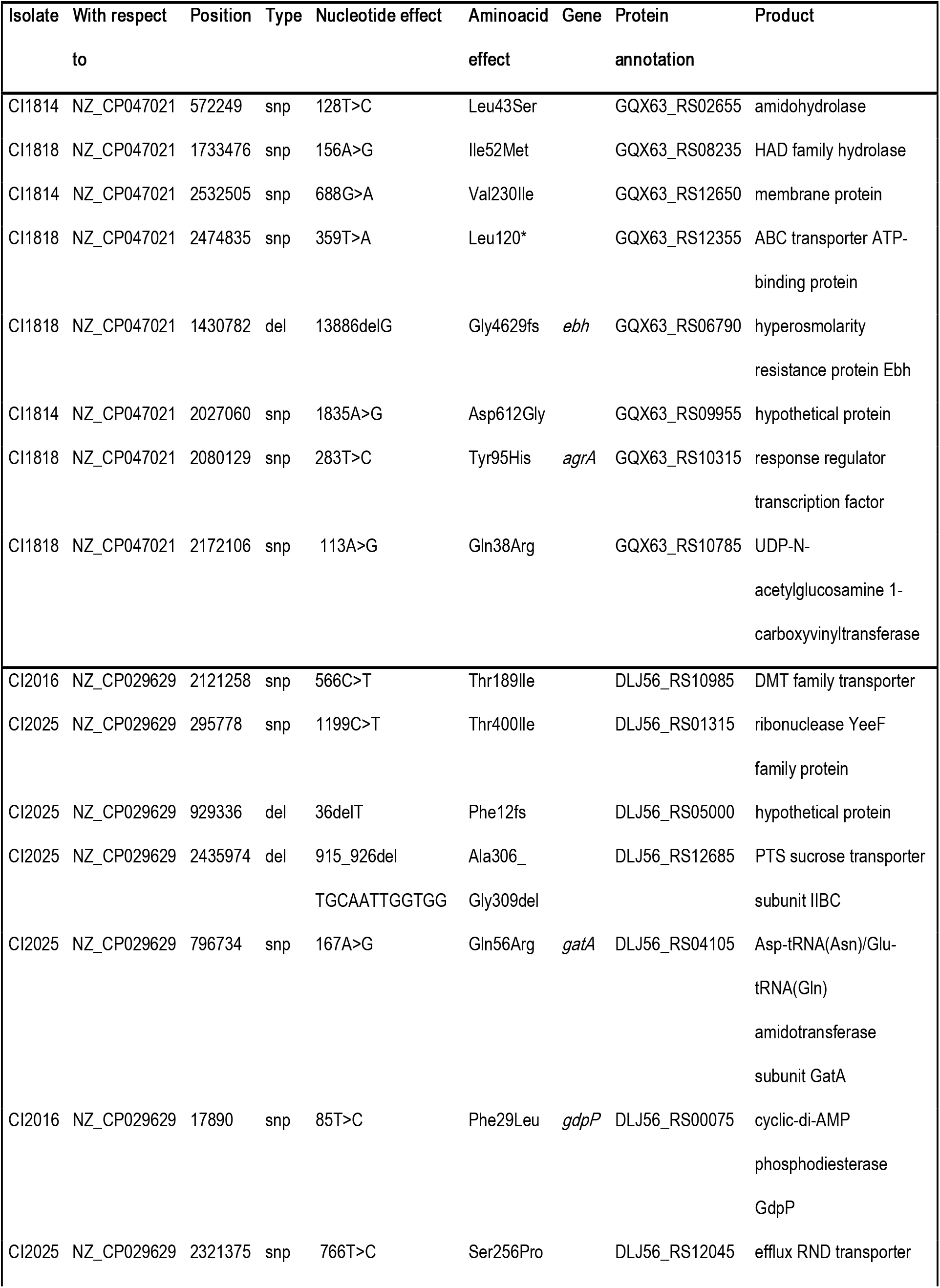
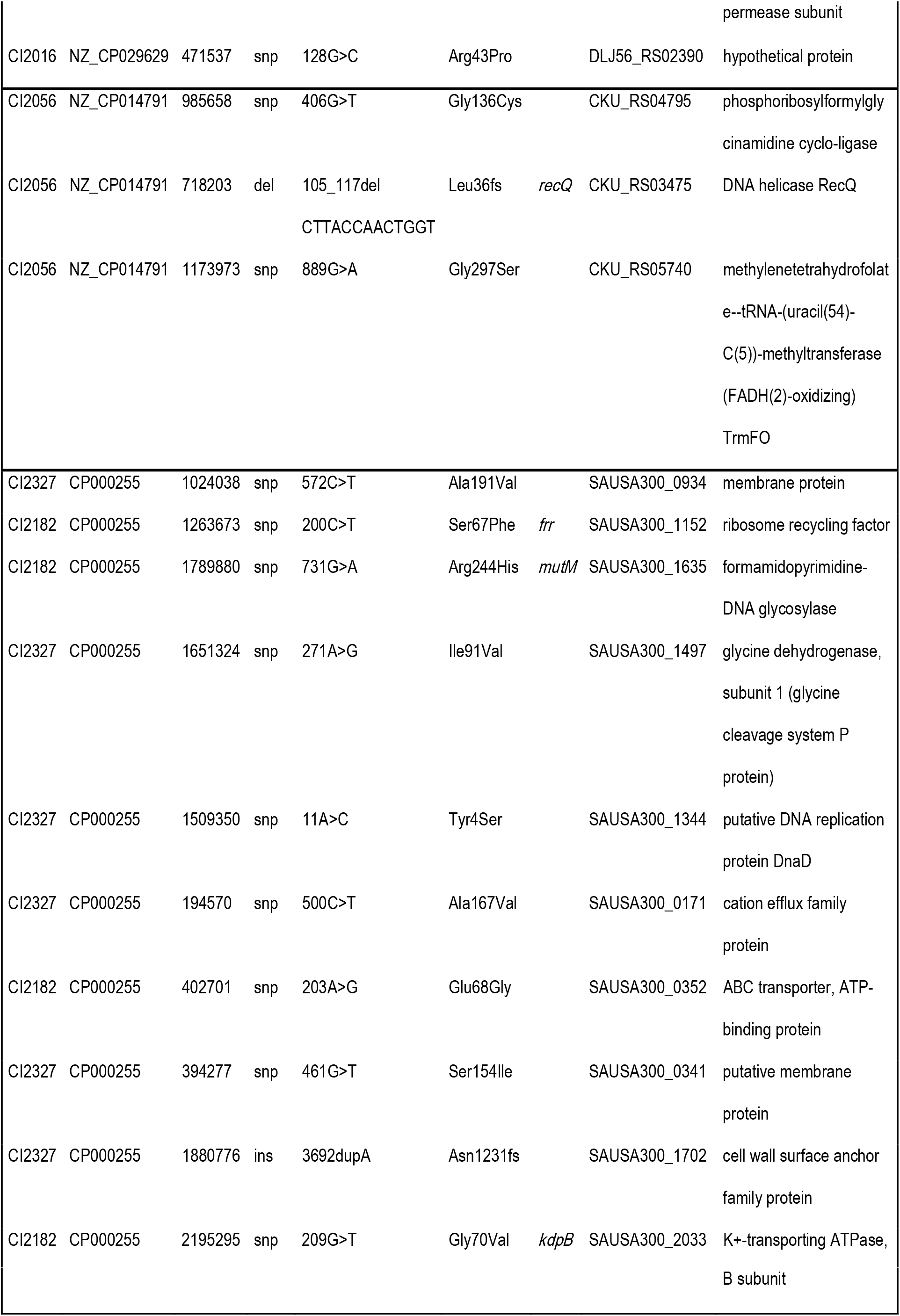
Non-synonymous mutations differentiating isolates’ pairs. This is a subset of Supplementary Table 3, which includes all mutations differentiating isolate pairs. Here only non-synonymous mutations are shown, as they might affect protein functions, and thus have a biological impact. The changes are given with respect to publicly available assembled references, whose GenBank accession are provided in the second column. The base-pair position, mutation type and effect at the nucleotide/amino-acid level, as well as the annotation and product of the affected proteins in the corresponding reference genome are shown. For each pair, a closely related reference was chosen. Importantly, note that the original allele/protein variant of the common ancestor of the two isolates is unknown. snp=single nucleotide polymorphism, del=deletion, ins=insertion, *=premature stop codon, fs=frameshift, nt=nucleotide, aa=amino acid

## References

1. Lowy FD. 1998. Staphylococcus aureus infections. N Engl J Med 339:520–32.

2. Kluytmans JA, Wertheim HF. 2005. Nasal carriage of Staphylococcus aureus and prevention of nosocomial infections. Infection 33:3–8.

3. Perl TM, Cullen JJ, Wenzel RP, Zimmerman MB, Pfaller MA, Sheppard D, Twombley J, French PP, Herwaldt LA. 2002. Intranasal mupirocin to prevent postoperative Staphylococcus aureus infections. N Engl J Med 346:1871–7.

4. Brugger SD, Bomar L, Lemon KP. 2016. Commensal-Pathogen Interactions along the Human Nasal Passages. PLoS Pathog 12:e1005633.

5. Casadevall A, Pirofski LA. 1999. Host-pathogen interactions: redefining the basic concepts of virulence and pathogenicity. Infect Immun 67:3703–13.

6. Laupland KB, Ross T, Gregson DB. 2008. Staphylococcus aureus bloodstream infections: risk factors, outcomes, and the influence of methicillin resistance in Calgary, Canada, 2000-2006. J Infect Dis 198:336–43.

7. Huemer M, Mairpady Shambat S, Brugger SD, Zinkernagel AS. 2020. Antibiotic resistance and persistence—Implications for human health and treatment perspectives. EMBO reports 21:e51034.

8. van Rijen MM, Bonten M, Wenzel RP, Kluytmans JA. 2008. Intranasal mupirocin for reduction of Staphylococcus aureus infections in surgical patients with nasal carriage: a systematic review. J Antimicrob Chemother 61:254–61.

9. Kluytmans J, van Belkum A, Verbrugh H. 1997. Nasal carriage of Staphylococcus aureus: epidemiology, underlying mechanisms, and associated risks. Clin Microbiol Rev 10:505–20.

10. Marshall C, McBryde E. 2014. The role of Staphylococcus aureus carriage in the pathogenesis of bloodstream infection. BMC Res Notes 7:428.

11. von Eiff C, Becker K, Machka K, Stammer H, Peters G. 2001. Nasal carriage as a source of Staphylococcus aureus bacteremia. Study Group. N Engl J Med 344:11–6.

12. Young BC, Wu CH, Gordon NC, Cole K, Price JR, Liu E, Sheppard AE, Perera S, Charlesworth J, Golubchik T, Iqbal Z, Bowden R, Massey RC, Paul J, Crook DW, Peto TE, Walker AS, Llewelyn MJ, Wyllie DH, Wilson DJ. 2017. Severe infections emerge from commensal bacteria by adaptive evolution. Elife 6.

13. Giulieri SG, Baines SL, Guerillot R, Seemann T, Gonçalves da Silva A, Schultz M, Massey RC, Holmes NE, Stinear TP, Howden BP. 2018. Genomic exploration of sequential clinical isolates reveals a distinctive molecular signature of persistent Staphylococcus aureus bacteraemia. Genome Med 10:65.

14. O’Donnell S, Humphreys H, Hughes D. 2008. Distribution of virulence genes among colonising and invasive isolates of methicillin-resistant Staphylococcus aureus. Clin Microbiol Infect 14:625–6.

15. Jenkins A, Diep BA, Mai TT, Vo NH, Warrener P, Suzich J, Stover CK, Sellman BR. 2015. Differential Expression and Roles of &lt;span class=&quot;named-content genus-species&quot; id=&quot;named-content-1&quot;&gt;Staphylococcus aureus&lt;/span&gt; Virulence Determinants during Colonization and Disease. mBio 6:e02272–14.

16. Tuchscherr L, Pöllath C, Siegmund A, Deinhardt-Emmer S, Hoerr V, Svensson CM, Thilo Figge M, Monecke S, Löffler B. 2019. Clinical S. aureus Isolates Vary in Their Virulence to Promote Adaptation to the Host. Toxins (Basel) 11.

17. Abu Othman A, Humphreys H, O’Neill E, Fitzgerald-Hughes D. 2011. Differences in expression of virulence genes amongst invasive and colonizing isolates of meticillin-resistant Staphylococcus aureus. J Med Microbiol 60:259–61.

18. EUCAST. 25 January 2020. http://www.eucast.org/clinical_breakpoints/. Accessed

19. Seidl K, Leimer N, Palheiros Marques M, Furrer A, Holzmann-Bürgel A, Senn G, Zbinden R, Zinkernagel AS. 2015. Clonality and antimicrobial susceptibility of methicillin-resistant Staphylococcus aureus at the University Hospital Zurich, Switzerland between 2012 and 2014. Ann Clin Microbiol Antimicrob 14:14.

20. Bolger AM, Lohse M, Usadel B. 2014. Trimmomatic: a flexible trimmer for Illumina sequence data. Bioinformatics 30:2114–20.

21. Bankevich A, Nurk S, Antipov D, Gurevich AA, Dvorkin M, Kulikov AS, Lesin VM, Nikolenko SI, Pham S, Prjibelski AD, Pyshkin AV, Sirotkin AV, Vyahhi N, Tesler G, Alekseyev MA, Pevzner PA. 2012. SPAdes: a new genome assembly algorithm and its applications to single-cell sequencing. J Comput Biol 19:455–77.

22. PubMLST. https://pubmlst.org/. Accessed

23. GitHub.

24. Seemann T. 2014. Prokka: rapid prokaryotic genome annotation. Bioinformatics 30:2068–9.

25. Page AJ, Cummins CA, Hunt M, Wong VK, Reuter S, Holden MT, Fookes M, Falush D, Keane JA, Parkhill J. 2015. Roary: rapid large-scale prokaryote pan genome analysis. Bioinformatics 31:3691–3.

26. Price MN, Dehal PS, Arkin AP. 2010. FastTree 2--approximately maximum-likelihood trees for large alignments. PLoS One 5:e9490.

27. Seemann T. 2020. Rapid haploid variant calling and core genome alignment.

28. Dengler Haunreiter V, Boumasmoud M, Haffner N, Wipfli D, Leimer N, Rachmuhl C, Kuhnert D, Achermann Y, Zbinden R, Benussi S, Vulin C, Zinkernagel AS. 2019. In-host evolution of Staphylococcus epidermidis in a pacemaker-associated endocarditis resulting in increased antibiotic tolerance. Nat Commun 10:1149.

29. Andreoni F, Ogawa T, Ogawa M, Madon J, Uchiyama S, Schuepbach RA, Zinkernagel AS. 2014. The IL-8 protease SpyCEP is detrimental for Group A Streptococcus host-cells interaction and biofilm formation. Front Microbiol 5:339.

30. Frey PM, Baer J, Bergadà Pijuan J, Lawless C, Bühler PK, Kouyos RD, Lemon KP, Zinkernagel AS, Brugger SD. 2020. Quantifying variation in bacterial reproductive fitness with and without antimicrobial resistance: a high-throughput method. bioRxiv doi:10.1101/2020.05.13.093807:2020.05.13.093807.

31. Wiegand I, Hilpert K, Hancock RE. 2008. Agar and broth dilution methods to determine the minimal inhibitory concentration (MIC) of antimicrobial substances. Nat Protoc 3:163–75.

32. Vulin C, Leimer N, Huemer M, Ackermann M, Zinkernagel AS. 2018. Prolonged bacterial lag time results in small colony variants that represent a sub-population of persisters. Nat Commun 9:4074.

33. Huemer M, Mairpady Shambat S, Bergada-Pijuan J, Söderholm S, Boumasmoud M, Vulin C, Gómez-Mejia A, Antelo Varela M, Tripathi V, Götschi S, Marques Maggio E, Hasse B, Brugger SD, Bumann D, Schuepbach RA, Zinkernagel AS. 2021. Molecular reprogramming and phenotype switching in Staphylococcus aureus lead to high antibiotic persistence and affect therapy success. Proc Natl Acad Sci U S A 118.

34. Bär J, Boumasmoud M, Kouyos RD, Zinkernagel AS, Vulin C. 2020. Efficient microbial colony growth dynamics quantification with ColTapp, an automated image analysis application. Sci Rep 10:16084.

35. Peleg AY, Monga D, Pillai S, Mylonakis E, Moellering RC, Jr., Eliopoulos GM. 2009. Reduced susceptibility to vancomycin influences pathogenicity in Staphylococcus aureus infection. J Infect Dis 199:532–6.

36. McCarthy AJ, Lindsay JA. 2010. Genetic variation in Staphylococcus aureus surface and immune evasion genes is lineage associated: implications for vaccine design and host-pathogen interactions. BMC Microbiol 10:173.

37. Christner M, Franke GC, Schommer NN, Wendt U, Wegert K, Pehle P, Kroll G, Schulze C, Buck F, Mack D, Aepfelbacher M, Rohde H. 2010. The giant extracellular matrix-binding protein of Staphylococcus epidermidis mediates biofilm accumulation and attachment to fibronectin. Mol Microbiol 75:187–207.

38. Clarke SR, Harris LG, Richards RG, Foster SJ. 2002. Analysis of Ebh, a 1.1-megadalton cell wall-associated fibronectin-binding protein of Staphylococcus aureus. Infect Immun 70:6680–7.

39. Cheng AG, Missiakas D, Schneewind O. 2014. The Giant Protein Ebh Is a Determinant of &lt;span class=&quot;named-content genus-species&quot; id=&quot;named-content-1&quot;&gt;Staphylococcus aureus&lt;/span&gt; Cell Size and Complement Resistance. Journal of Bacteriology 196:971.

40. Thomsen IP, Kadari P, Soper NR, Riddell S, Kiska D, Creech CB, Shaw J. 2019. Molecular Epidemiology of Invasive Staphylococcus aureus Infections and Concordance with Colonization Isolates. J Pediatr 210:173–177.

41. Berube BJ, Bubeck Wardenburg J. 2013. Staphylococcus aureus alpha-toxin: nearly a century of intrigue. Toxins (Basel) 5:1140–66.

42. Tam K, Torres VJ. 2019. Staphylococcus aureus Secreted Toxins and Extracellular Enzymes. Microbiol Spectr 7.

43. Sinha B, Fraunholz M. 2010. Staphylococcus aureus host cell invasion and post-invasion events. Int J Med Microbiol 300:170–5.

44. Foster TJ, Geoghegan JA, Ganesh VK, Höök M. 2014. Adhesion, invasion and evasion: the many functions of the surface proteins of Staphylococcus aureus. Nature reviews Microbiology 12:49–62.

45. Huemer M, Mairpady Shambat S, Bergada-Pijuan J, Söderholm S, Boumasmoud M, Vulin C, Gómez-Mejia A, Antelo Varela M, Tripathi V, Götschi S, Marques Maggio E, Hasse B, Brugger SD, Bumann D, Schuepbach RA, Zinkernagel AS. 2021. Molecular reprogramming and phenotype switching in &lt;em&gt;Staphylococcus aureus&lt;/em&gt; lead to high antibiotic persistence and affect therapy success. Proceedings of the National Academy of Sciences 118:e2014920118.

46. Sheehan G, Garvey A, Croke M, Kavanagh K. 2018. Innate humoral immune defences in mammals and insects: The same, with differences ? Virulence 9:1625–1639.

47. Lindsay JA, Moore CE, Day NP, Peacock SJ, Witney AA, Stabler RA, Husain SE, Butcher PD, Hinds J. 2006. Microarrays reveal that each of the ten dominant lineages of Staphylococcus aureus has a unique combination of surface-associated and regulatory genes. Journal of bacteriology 188:669–676.

48. Deinhardt-Emmer S, Sachse S, Geraci J, Fischer C, Kwetkat A, Dawczynski K, Tuchscherr L, Loffler B. 2018. Virulence patterns of Staphylococcus aureus strains from nasopharyngeal colonization. J Hosp Infect 100:309–315.

49. Richards RL, Haigh RD, Pascoe B, Sheppard SK, Price F, Jenkins D, Rajakumar K, Morrissey JA. 2015. Persistent Staphylococcus aureus isolates from two independent cases of bacteremia display increased bacterial fitness and novel immune evasion phenotypes. Infection and immunity 83:3311–3324.

50. Jenkins A, Diep BA, Mai TT, Vo NH, Warrener P, Suzich J, Stover CK, Sellman BR. 2015. Differential expression and roles of Staphylococcus aureus virulence determinants during colonization and disease. mBio 6:e02272.

51. Camargo ILBC, Gilmore MS. 2008. Staphylococcus aureus--probing for host weakness? Journal of bacteriology 190:2253–2256.

